# DNA extraction and virome processing methods strongly influence recovered human gut viral community characteristics

**DOI:** 10.1101/2025.11.25.690293

**Authors:** Luke S. Hillary, Trina A. Knotts, Sean H. Adams, Mohamed R. Ali, Matthew R. Olm, Joanne B. Emerson

## Abstract

Accurately characterising the human gut virome is critical to understanding virus-microbiome-host interactions. However, widely used methods introduce biases that complicate data interpretation and limit cross-study comparability. For instance, multiple-displacement amplification (MDA) preferentially amplifies single-stranded DNA viruses, while total metagenomes are dominated by non-viral sequences, reducing viral signal. These traditional methods have not been systematically compared to viral size-fraction metagenomes (viromes) prepared without MDA. To address this, we applied four common methods for characterising human gut viral community composition (total metagenomes, viromes with/ without DNase treatment (to remove free DNA), and MDA viromes) to a human stool sample, with technical triplicates for each approach. MDA biased viral community composition to a shocking degree: *Microviridae* formed ∼90% of MDA viromes compared to just 2% of non-MDA viromes. Removing ssDNA viruses from data analyses substantially reduced, but did not eliminate, MDA bias. Metagenomes were enriched for putative temperate phages and predicted *Bacillota-phages*, whereas predicted *Bacteroidetes*-phages dominated all viromes, suggesting that metagenomes and viromes select for different populations within the total viral community. DNase treatment had little-to-no effect on virome richness or community composition. This proof-of-principle experiment demonstrates that preparatory methods for viral community analysis can lead to substantially different conclusions from the same faecal sample, and we provide a comprehensive omic data analysis framework for comparing laboratory methodologies for viral ecology. With sufficient DNA yields now easily achievable from human gut viromes without the use of MDA, our results suggest that this biased amplification method should be avoided in human gut virome studies.

Graphical Abstract

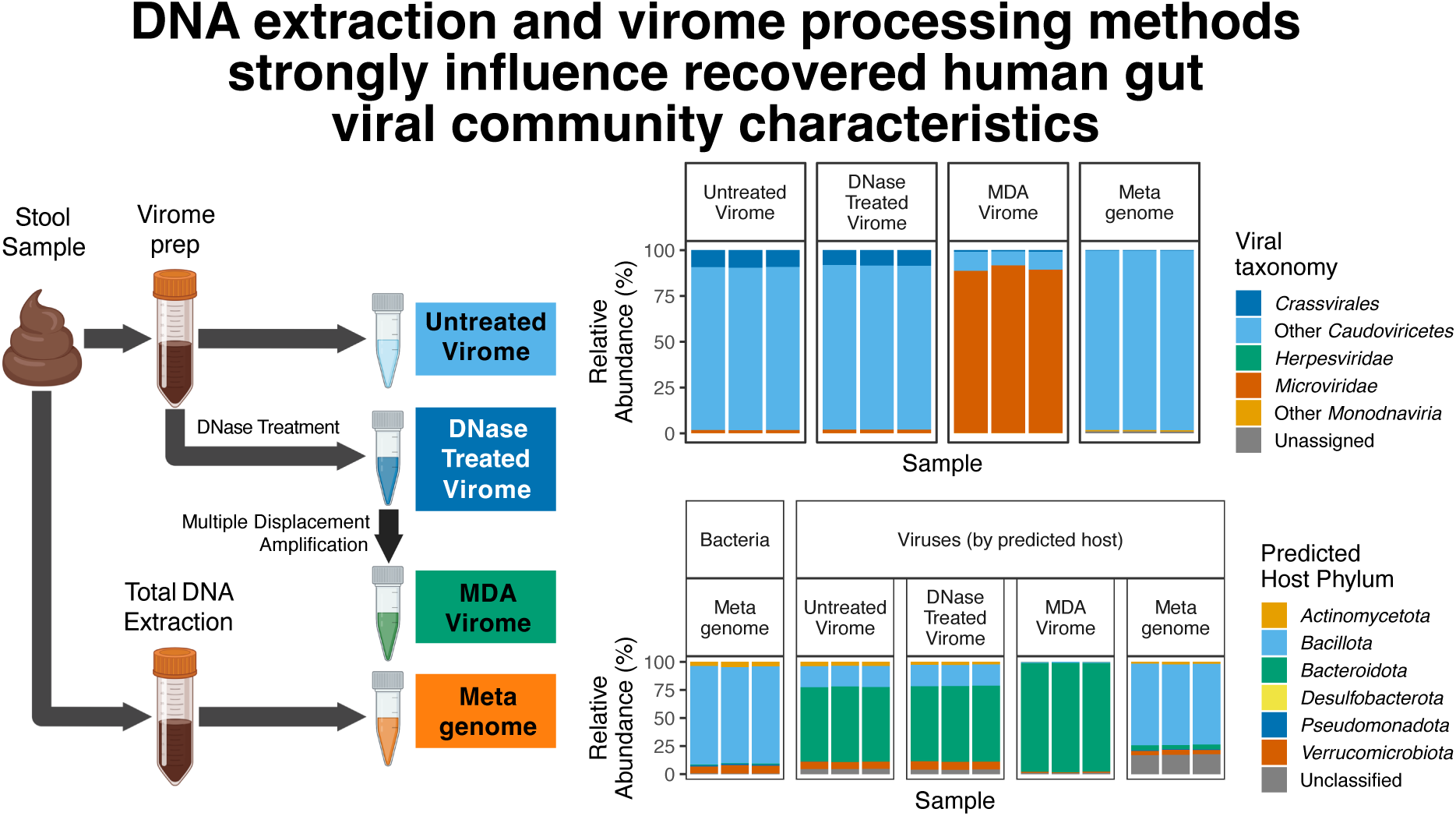

## Introduction

The viruses present in the human gastrointestinal tract, otherwise known as the gut virome, have the potential to regulate the composition, activity, and functional capacity of the gut microbiome through cell lysis, altered cellular metabolism, and horizontal gene transfer (1,2). Viral dysbiosis, a disruptive imbalance of the human gut virome (e.g. lower diversity), is associated with various disease states and may act as a diagnostic indicator in inflammatory bowel disease and pancreatic cancer (3–6). Phage therapy, via individual phages, phage cocktails, and whole-virome transplants, has the potential to address issues related to antimicrobial resistance and targeted interventions in gastrointestinal diseases (7,8). Therefore, accurate characterisation of the gut virome composition and dynamics is of critical importance.

Human gut virome studies often use either faecal shotgun metagenomics or size-selective viromics to characterise viral community composition, functional capacity, and ecological dynamics (9–11). Comparative studies in soils and aquatic environments show that while total metagenomes capture integrated prophages, giant viruses excluded by size-selection, and actively-replicating viruses often absent from the viral size fraction, they can recover less fine-scale diversity than size-selective viromes (12–14). Thus, metagenomics and viromics can differ in recovered viral community compositions (15), but systematic comparisons across different techniques remain limited.

Previous work on improving viral community characterization has focused on optimizing sample handling (16), viral concentration methods (17,18), nucleic acid extraction (19,20), PCR amplification cycles (21), and sequencing library preparation (22). As many of these studies focused on mock communities or viral spike-ins, viral recovery and diversity metrics were the primary evaluation criteria. This leaves clear knowledge gaps in how different methods affect recoverable virome diversity within complex human gut communities.

The amount of faecal material available for DNA extraction can be limited, leading to an assumed need for amplification to achieve sufficient yields for sequencing. This is often accomplished via high numbers of PCR cycles (>6-8) during sequencing library preparation or through an initial treatment of multiple displacement amplification (MDA) before library construction (21,23). However, MDA is known to bias viromes towards amplifying small circular DNA, leading to the presumed but rarely quantified overrepresentation of single-stranded DNA viruses, such as *Microviridae*, in MDA-viromes from diverse environments (24,25). Given this known bias, systematic tests comparing viral communities recovered from MDA viromes and non-MDA viromes are needed.

Although the effects of DNase treatment have not been systematically evaluated for human gut viromes, studies in soil show that freezing can lead to viromic DNA yields below detection limits after DNase treatment (26,27), presumably because freezing compromises virions and exposes viral genomic DNA to enzymatic degradation. In agricultural soils, DNase treatment on fresh samples resulted in a 53% reduction in viral diversity, although broader community structure and ecological patterns remained similar with and without DNase treatment (28). As most faecal samples from human studies are stored frozen for logistical reasons, and storage above freezing has been shown to alter microbial community composition (29), the effects of DNase treatment on previously-stored faecal viromes warrant further consideration. Given that omitting DNase treatment improves viral recovery from frozen soil samples, the potential for skipping DNase treatment on faecal samples stored frozen should be evaluated, especially since low DNA yields might make biased MDA more tempting.

Here we leveraged our soil viromics protocol (largely similar to existing protocols in the human gut literature - Conceição-Neto et al., 2015; Kleiner et al., 2015; Wang et al., 2023) to: (i) compare two types of virome preparations (with and without DNase treatment – Sorensen et al., 2021), (ii) add a multiple-displacement amplification (MDA) treatment to aliquots from the DNase-treated viromes, an approach commonly applied in the human gut virome literature (31), and (iii) compare all of the virome preparations to “total metagenomes,” commonly used for viral community recovery from a variety of ecosystems, including soil and the human gut (32,33). With analyses considering percent viral reads, viral contigs assembled *de novo* vs. recoverable through read mapping to a reference set, viral genome quality, viral richness, viral community beta-diversity, viral taxonomy, predicted host taxonomy, predicted viral replication strategies, library k-mer complexity, functional annotations, viral genome coverage depth to enable strain-level analyses, and comparisons to a comprehensive gut viral genomic database, we also provide an example for rigorous comparisons of the downstream omic data resulting from methodological differences. Since the faecal material here was stored frozen (experiencing a single freeze-thaw), we hypothesised that DNase treatment would yield insufficient DNA for sequencing without further amplification and that viromes without DNase treatment would recover the highest viral diversity. We hypothesised that these viromes would recover vastly more viral species than total metagenomes and that MDA viromes would preferentially recover *Microviridae* (small, single-stranded circular DNA viruses), rendering MDA viromes non-quantitative and not representative of the non-MDA viral community. To test these hypotheses and provide guidance for future studies, we systematically compared untreated, DNase-treated, and MDA viromes against metagenomes from the same single faecal source material. This design allowed us to directly assess how upstream processing choices shape downstream characterisation of the gut virome, providing an observational benchmark to inform future experimental design.

## Materials and methods

### Sample collection and study design

Faecal sub-samples were generated from a single stool sample collected from a healthy female participant recruited from the UC Davis bariatric surgery clinic during the pre-operative period, and while consuming a typical diet. The participant provided informed consent, sample collection protocols were approved by the UC Davis Institutional Review Board, and the studies conform to the Declaration of Helsinki (34). The stool material was stored intact at −80 °C until processing. A 100 g stool sample was thawed overnight at 4°C and manually homogenised, followed by sub-sampling for DNA extraction.

In total, four approaches to measure viral community composition were compared in triplicate (see Supplementary Fig. S1): untreated viromes, DNase-treated viromes, MDA viromes (DNase-treated viromic DNA that was subsequently treated with multiple-displacement amplification, MDA), and stool total metagenomes.

### Virome sample processing

For viromes, virus-like particles (VLPs) were purified from six 10 g sub-samples of thawed stool as previously described for soil (35). Briefly, VLPs were purified from stool by three sequential rounds of suspension in 9 mL of protein-supplemented PBS buffer (PPBS - 2% bovine serum albumin, 10% phosphate-buffered saline, 150 mM MgSO_4_) in 50 mL conical tubes, each followed by 10 minutes of orbital shaking at 300 rpm, 4°C, and 10 minutes centrifugation at 4,000 × g, 4 °C. Supernatants from the first and second rounds were stored in separate 50 mL conical tubes at 4 °C during subsequent rounds of resuspension, shaking, and centrifugation. Supernatants from each aliquot were combined, centrifuged at 10,000 × g at 4°C for 8 minutes, and the supernatant was retained, followed by a second round of centrifugation. Supernatants were filtered sequentially through 5 µm, 0.45 µm, and 0.2 µm PES (polyethersulfone) syringe filters and pooled in 50 mL conical tubes prior to being transferred to 26.3 mL polycarbonate round-bottomed ultracentrifuge tubes (Beckman-Coulter Life Sciences), followed by ultracentrifugation at 35,000 rpm (112,000 × g) at 4 °C for 145 minutes using an Optima LE-80K ultracentrifuge and 50.2 Ti rotor (Beckman-Coulter Life Sciences). Supernatants were discarded, and pellets resuspended in 600 µL nuclease-free water. DNase treatment was performed on three 100 µL aliquots of VLP concentrate by incubating them with 10 µL RQ1 DNase and 10 µL RQ1 DNase buffer (Promega) for 30 minutes at 37 °C. DNase was inactivated by the addition of 10 µL of RQ1 DNase stop solution (Promega).

### Virome and metagenome DNA extraction, multiple-displacement amplification, and library construction

DNA was extracted from 100 µL aliquots of VLP concentrates (untreated viromes), the DNase-treated virome preparations, or 0.25 mg of stool (for metagenomes), using the PowerSoil Pro DNA extraction kit (Qiagen) per manufacturer’s instructions. Samples were incubated with lysis buffer for 10 minutes at 65 °C followed by vortexing for 10 minutes at maximum speed. Further steps were carried out according to the manufacturer’s instructions. DNA was quantified using a Qubit 1x High Sensitivity DNA quantification assay and Qubit 4 fluorimeter (Thermo Fisher Scientific, Inc.).

To generate the MDA viromes, three 1 µL aliquots of DNA from each of the three DNase-treated virome preparations were used as templates for MDA, for nine reactions in total. MDA was performed using the GenomiPhi V2 DNA amplification kit (Cytiva), according to the manufacturer’s instructions. The nine reactions were then pooled into three treatment replicates (three MDA reactions per DNase-treated virome), yielding three MDA virome preparations.

Libraries for all four methods (3× untreated viromes, 3× DNase-treated viromes, 3× MDA viromes, 3× metagenomes) were prepared using the KAPA DNA HyperPrep library kit (Roche) by the DNA Technologies & Expression Analysis Core, UC Davis, and 150 bp paired-end sequencing was performed to a target depth of 20 Gbp per library using the Illumina NovaSeq 6000 platform.

### Sequencing data quality control

Detailed data processing settings are provided in Supplementary Table S2. Raw reads were trimmed and filtered using BBDuk v39.1, error-corrected using Tadpole, and deduplicated using Clumpify (all part of BBTools v39.1 Bushnell 2018). Raw and processed read quality was assessed using FastQC v0.12.1 (Andrews 2010) and MultiQC v1.14 (Ewels et al. 2016).

### Library complexity, rRNA gene read, and human read analyses

K-mer (k=31) frequencies were calculated using khmer v2.1.1 (36). Ribosomal RNA gene reads were identified using SortMeRNA v4.3.6 (37). Taxonomic profiling of raw reads was performed using SingleM v0.18.3 (38). To identify human host reads, error-corrected reads were mapped to the human genome (GCF_009914755.1, Nurk et al., 2022) using Minimap2 v2.26 (40) and Samtools v1.17 (41). Count data were aggregated using CoverM v0.6.1 (42).

### Assembly and viral contig identification

Reads from each library were separately assembled (12 assemblies) using MEGAHIT v1.2.9 (43), and summary statistics were produced using Quast v5.2.0 (44). Viral contigs were identified from each assembly using geNomad v1.7.0 (45) and additionally filtered by length ≥10 Kbp (46) or with a requirement for both a length between 1 and 10 Kbp and a geNomad phylum = *Monodnaviria*. The latter requirement facilitates inclusion of single-stranded DNA viruses that typically possess genomes <10 Kbp (47).

### vOTU clustering, read mapping, and community compositional analyses

All viral contigs were clustered together into vOTUs, using a combination of MegaBLAST v2.14.0 and custom Python scripts (see Data and Code Availability), requiring a minimum average nucleotide identity (ANI) of 95% across 85% of the length of the shortest contig (48–50). Quality-filtered and error-corrected reads from each library were mapped to dereplicated vOTUs using minimap2 (40) and relative abundances calculated using transcripts per million (TPM) values generated by CoverM v0.6.1 (42).

The presence of vOTUs in a sample was determined using two complementary criteria: detection by read mapping and recovery of viral contigs through a combination of mapping and assembly. For vOTUs categorised as “mapped” a vOTU was considered shared between two libraries if ≥75% of its length was covered at ≥ 1x read depth and 90% ANI in both libraries. vOTUs were considered “assembled” in a library if, in addition to reads mapping to the vOTU sequence according to the aforementioned thresholds, a viral contig from the same vOTU cluster was assembled from that same library.

To identify known vOTUs from the Unified Human Gut Virome catalogue (UHGV, https://github.com/snayfach/UHGV, accessed on June 12^th^ 2025) (31,49,51–60) within libraries from this study, all UHGV vOTUs >10Kbp in length or >50% complete were downloaded from https://portal.nersc.gov/UHGV/ on June 12^th^ 2025. These UHGV vOTUs were clustered with all 2,480 predicted viral contigs from this study using the same parameters as above, and reads from our libraries were mapped to this dereplicated set. A UHGV vOTU was considered detected in our dataset if it clustered with at least one viral sequence from this study or if read mapping covered the vOTU in at least one sample at the same detection thresholds described above.

### Read depth analysis

Raw reads were randomly sub-sampled to depths of 1 Gbp increments from 1-17 Gbp using seqtk v1.4-r122 (https://github.com/lh3/seqtk), and quality control, assembly, viral contig identification, vOTU clustering, and read mapping were performed on each subsampled dataset as described above.

### Viral translated protein annotation

As the gene content of vOTU cluster sequences can differ from the vOTU representative sequence, all viral translated proteins from predicted viral contigs by geNomad were annotated in bulk using Pharokka v1.7.1 with database v1.4.0 (61).

### vOTU lifestyle and host predictions

Viral lifestyle, i.e., whether a virus is virulent or temperate, was predicted using BACPHLIP v0.9.6 (62). vOTUs were classified as putatively virulent or temperate if the confidence score for either assignment was ≥0.95; otherwise, they were labelled unclassified. vOTU host prediction was performed using iPHoP v1.3.2, with a combined prediction confidence score to genus level of ≥90/100 (63). When two hosts were predicted, the host with higher confidence was selected. If confidence scores were equal, host prediction was converted to the lowest common taxonomic level.

### Data analysis and visualization

Statistical analyses and data visualization were performed using R v4.4.0, RStudio v2024.09.1-394, ggpubr (64), and tidyverse (65). Statistically significant differences with p-values <0.05 were identified using analysis of variance (ANOVA) and post-hoc Tukey tests and converted into compact letter displays using multcompView (66). Venn diagrams were visualised using ggVennDiagram (67). Principal Coordinates Analyses (PCoA) of pairwise Bray-Curtis dissimilarity matrices were performed using Vegan (68), and significant differences were identified by permutational multivariate analysis of variance (PERMANOVA).

## Results and discussion

### Recovered viral communities differed across all processing methods, except between DNase-treated and untreated viromes

To evaluate the impacts of total metagenomics, virus-like particle (VLP) fractionation (viromics), DNase treatment, and multiple displacement amplification (MDA) on recoverable faecal viral community composition (Supplementary Fig. S1a), 12 libraries were generated from one stool sample, sequenced to an average depth of 21.1 ± 2.3 Gbp, and analysed. A total of 2,480 viral contig sequences were identified using geNomad (45) and clustered into 605 viral operational taxonomic units (vOTUs; Supplementary Fig. S2). Full results of statistical tests (ANOVA and Tukey post hoc comparisons) for all analyses are provided in Supplementary Tables S5–S6.

Because all preparations were performed on aliquots of the same faecal sample, observed differences reflect the influence of processing method rather than inter-individual variation, which is a known major source of variability in gut virome studies (2). While this single-sample design limits generalisability, it provides a clear view of methodological biases in isolation, supporting observational conclusions about their effects on detectable viral community composition and downstream analyses typically employed in intervention-based human gut virome studies.

The similarity between DNase-treated and untreated viromes following frozen stool sample storage was unexpected, as DNase treatment is often discouraged for environmental samples stored frozen since it can result in a 10-to 100-fold loss in viromic DNA yield (69). The negligible impact observed here likely reflects stool-specific factors, such as (i) higher viral loads and organic content, supporting higher intact viral particle recovery (15) and/or new virus production during thawing or (ii) a naturally high ratio of encapsidated to free DNA due to DNase I secretion in the small intestine (70). Unlike soils, where freeze-thaw cycles may damage a greater proportion of viral capsids (potentially due to higher viral diversity and overall lower biomass), stool VLP concentrates appear to retain sufficient encapsidated viral DNA to permit effective sequencing of frozen samples, even after DNase treatment. Together, these findings are particularly relevant to studies using frozen, biobanked, or transported samples without guaranteed cold-chain preservation above 0 °C. Of course, further tests using additional samples and comparisons between fresh and frozen stool samples would be required to assess the generalizability of these results. As DNase-treated and untreated viromes were virtually indistinguishable in each of our analyses in this study, we generally describe their results together and refer to them collectively as non-MDA viromes, but we retain them separately in calculations and figures.

### Method-dependent impacts on vOTU recovery, proportions of viral and rRNA gene reads, and viral community profiling

To evaluate the effects of virome and metagenome preparation methods on vOTU recovery, we assessed vOTU detection using two metrics: “assembled” vOTUs or “mapped” vOTUs. Notably, vOTUs only detected via read mapping for a given processing method would not have been detected if other processing methods had not been performed to generate the reference set of 605 vOTUs, whereas “assembled” vOTUs would have been detected from that method alone (see Methods and Supplementary Fig. S3). Both metrics produced similar patterns across processing methods. DNase-treated and untreated viromes recovered highly overlapping vOTUs, with 97% shared by mapping, confirming highly similar compositions. In contrast, 50% of vOTUs assembled uniquely in MDA viromes, and 62% of vOTUs uniquely assembled in metagenomes were not detected by other methods, even after read mapping. Only 25% of vOTUs were detected in at least one replicate from all four methods via read mapping (Fig. 1a-b). Metagenomes and non-MDA viromes assembled similar numbers of vOTUs (292–304), approximately half of the total, while MDA viromes only assembled 166 vOTUs (27%). By both metrics, MDA viromes shared more vOTUs with non-MDA viromes than with metagenomes, while metagenomes yielded the most unique vOTUs. These findings indicate that metagenomes and viromes recover distinct portions of the viral community, and differences persist when using a shared reference database for read mapping.

**Figure 1.**
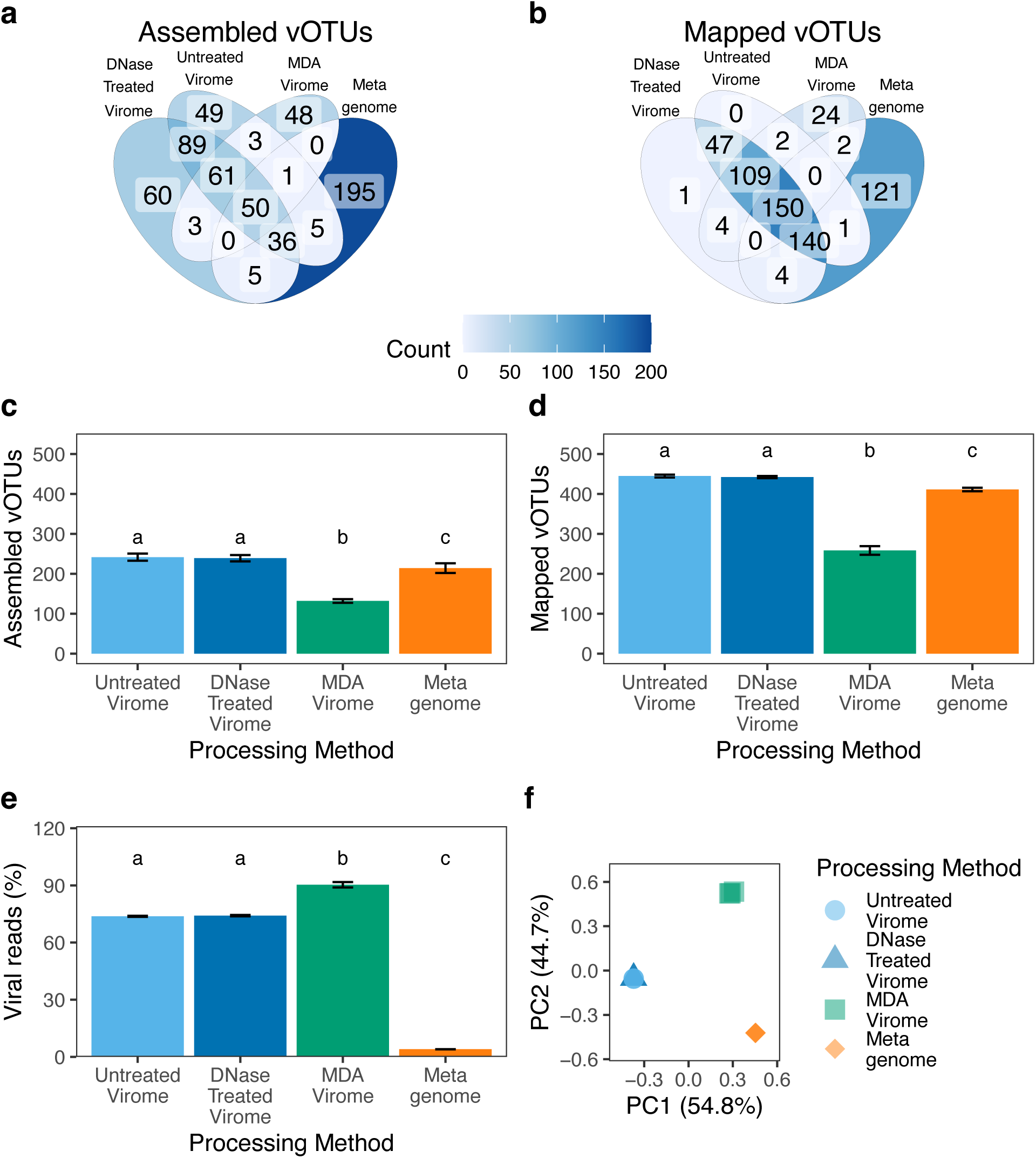
Comparison of vOTU recovery, read composition, and viral community structure across methods. (a-b) Venn diagrams showing the number of vOTUs shared between methods based on (a) assembly and (b) read mapping to a dereplicated set of 605 vOTUs. (c-d) Mean number of vOTUs detected per library in each method by (c) assembly and (d) read mapping. (e) Percentage of reads per library mapped to vOTUs by method. For panels (c-e): bars = mean, error bars = standard deviation, letters represent groupings according to Tukey HSD where differences require p_adjusted_<0.05. (f) Principal Coordinates Analysis (PCoA) of Bray-Curtis dissimilarities between samples with percent variance explained by PC1 and PC2. A total of 12 samples are shown, but due to the extreme similarity between some, differences among technical replicates are not visible (e.g., all three metagenome samples directly overlap). Significant differences among methods were tested by PERMANOVA, R^2^ = 0.996, p < 0.001.

Other alpha-diversity metrics support vOTU detection metrics, namely that non-MDA viromes, MDA viromes, and total metagenomes captured distinct parts of the viral community, with DNase-treated and untreated viromes statistically indistinguishable. On a per-library basis, both types of non-MDA viromes yielded significantly higher viral richness (number of vOTUs detected per library) than did MDA viromes or metagenomes by both “assembled” (Fig. 1c; ANOVA, F_3,8_=85.3, p < 00001; Tukey HSD, all p*_adjusted_* < 0.001) and “mapped” (Fig. 1d; ANOVA F_3,8_ = 31.5, p < 0.00001,; Tukey HSD, all p*_adjusted_* < 0.01 for non-MDA vs. MDA viromes) vOTU detection metrics. The MDA viromes had a mean of 132 assembled vOTUs, or only 55% of the vOTUs from the DNase-treated viromes from which the MDA virome DNA was sourced.

Despite this, MDA viromes had the highest proportion of viral reads (Fig. 1e; 91%), followed by non-MDA viromes (74% for both DNase-treated and untreated). As expected, metagenomes contained far fewer viral reads (4%; ANOVA, F_3,8_ = 1,720.8, p < 0.00001; Tukey p < 0.0001 for all viromes vs. metagenomes). These trends were mirrored in rRNA gene read proportions (a proxy for cellular organism-derived DNA), which were highest in the metagenomes (Supplementary Fig. S4b; ANOVA, F_3,8_ = 546.5, p < 0.00001). Human reads were slightly more abundant in the non-MDA viromes (0.22–0.25%) compared to the MDA viromes (0.14%) and metagenomes (0.12%; Supplementary Fig. S4c), but overall, these differences were not significant (ANOVA, F_3,8_ = 1.42, p=0.31). These results show that non-MDA viromes best recover viral diversity while minimizing cellular DNA contamination.

Beta-diversity patterns based on Bray-Curtis dissimilarities clearly separated the dataset into the three methodological groups. Principal Coordinates Analysis (PCoA, Fig. 1f) revealed strong separation into three clusters corresponding to non-MDA viromes, MDA viromes, and metagenomes (PERMANOVA, R² = 0.996, p < 0.001). Replicates clustered with minimal differences within methods, with DNase-treated and untreated viromes grouping together (ANOVA on distances to centroids F_1,4_ = 3.08, p = 0.091).

Within-method dissimilarities were consistently low (0.01-0.1), while between-group dissimilarities were substantially higher (0.88-0.96), except between DNase-treated and untreated viromes (0.04; Supplementary Fig. S5). These dissimilarity values are comparable to intra-subject or between-timepoint values from previous gut virome studies (71,72), suggesting that the processing method can overshadow biological signals in viral community composition.

### Multiple-displacement amplification severely biased viral taxonomic diversity, but the biases were largely computationally correctable

Despite known MDA biases, limited quantitative comparisons may contribute to the continued use of MDA viromes. To address this, we offer an empirical comparison of MDA viromes and non-MDA viromes. As expected, high-level taxonomic classification by geNomad (45) revealed that MDA viromes were overwhelmingly dominated by Microviridae, a family of small, circular, single-stranded DNA viruses, forming 90% relative abundance from 23-24 vOTUs per sample, compared to just 2% in non-MDA viromes (Fig. 2a). The staggering dominance of Microviridae in the MDA-viromes (45 times that of non-MDA viromes) bears emphasis here and adds tangible, quantitative, empirical rigor to previous suggestions of MDA bias. Conversely, the widespread dsDNA phage order Crassvirales (73), known to infect Bacteroidota and typically characterised by circular genomes of ∼100 kbp (alpha–gamma families) or ∼145–192 kbp (epsilon and zeta families) (74,75), was largely absent from MDA viromes (1%) and even more so from metagenomes (0.3%). In contrast, *Crassvirales* comprised 8-9% of the viral communities in non-MDA viromes, suggesting that MDA viromes may undersample this group. Since k-mer profiles reflect sequence diversity independently of taxonomic assignment (76–78), we also compared k-mer abundance distributions across methods to further assess how MDA affected sequence composition (Fig. 2b and Supplementary Fig. S4d). MDA viromes showed an underrepresentation of low-abundance k-mers compared to non-MDA viromes and metagenomes, consistent with lower sequence diversity in MDA viromes. Overall, MDA viromes were severely biased in recoverable viral taxonomic diversity compared to all other processing methods, leaving it difficult to justify using multiple-displacement amplification for viromics in the future if it can be avoided.

**Figure 2.**
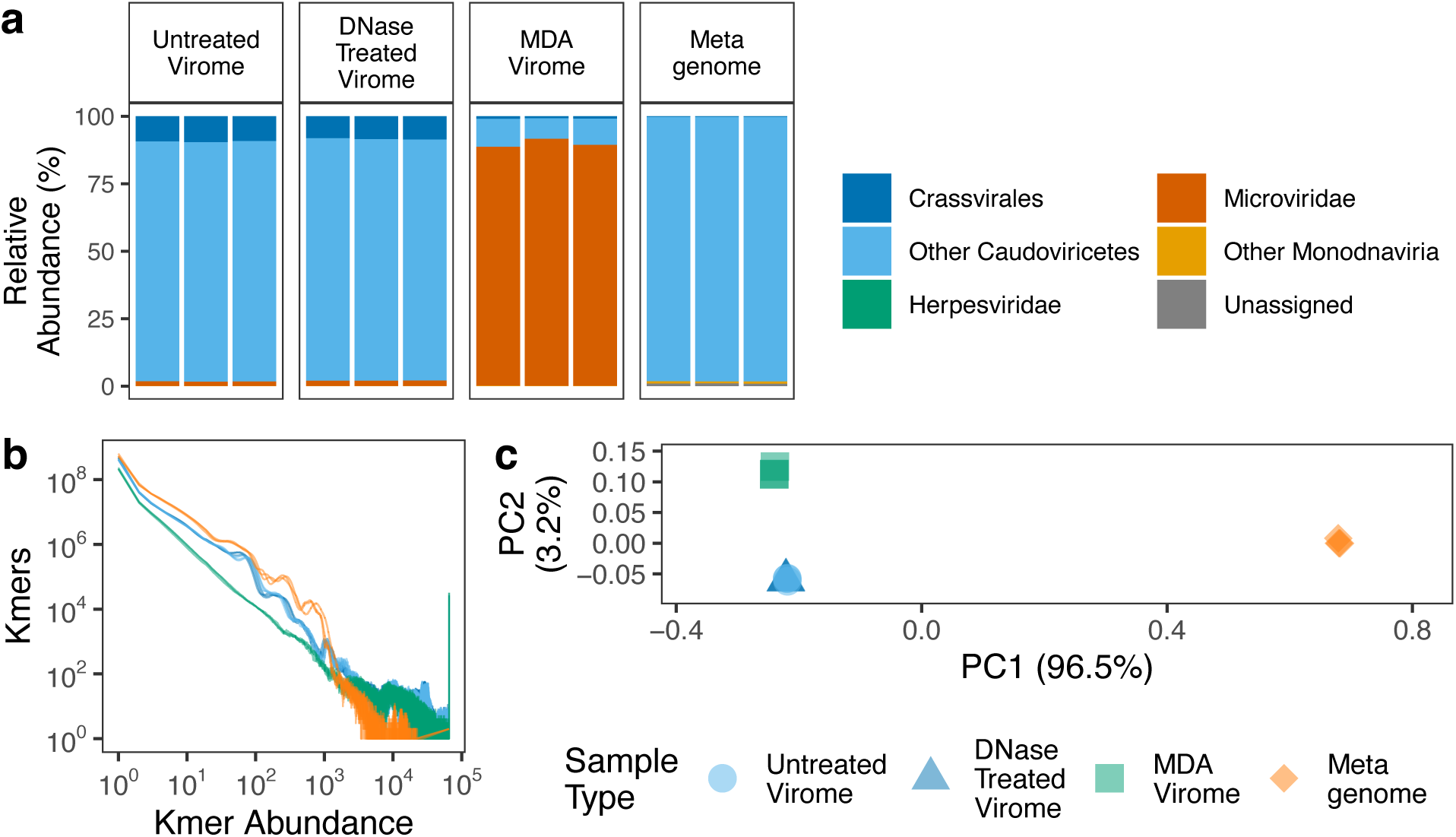
Impact of method on viral taxonomic composition and library k-mer profiles, and demonstrated potential to compensate for MDA compositional biases. (a) Relative abundances of viral taxa across methods, based on geNomad taxonomic classifications (45). (b) K-mer abundance profiles - MDA viromes in green show an underrepresentation of low-abundance k-mers (left side of the plot) compared to high abundance k-mers (right side of the plot). (c) Principal Coordinates Analysis (PCoA) of Bray-Curtis dissimilarities after removal of ssDNA viruses and recalculation of relative abundances. Methods remained significantly different but with 96.5% of the variance explained by PC1, corresponding to differences between viromes and metagenomes (PERMANOVA, 999 iterations, F = 1545.2, R^2^ = 0.998, p = 0.001).

To assess whether excluding ssDNA viruses mitigated MDA biases, we recalculated vOTU relative abundances after removing all ssDNA vOTUs, including the circular ssDNA families *Circoviridae, Genomoviridae, Inoviridae, and Microviridae*. No linear ssDNA viruses (*Spiraviridae, Bidnaviridae, Parvoviridae*) were detected. After filtering, 96.5% of the variance was attributable to differences between metagenomes and all viromes, regardless of DNase or MDA treatment (Fig. 2c). This increased observed compositional similarity across all viromes and separation of viromes from metagenomes after removing ssDNA viral data was confirmed by PERMANOVA (R^2^ = 0.998, p < 0.001), suggesting that MDA viromes may still retain useful biological signal after dominant ssDNA vOTUs are excluded.

These findings highlight the substantial taxonomic distortions introduced by MDA. Still, it is encouraging that some biases can be reduced computationally to potentially make existing datasets (or future datasets if MDA is unavoidable) more representative. While MDA bias towards small circular ssDNA viruses such as *Microviridae* is known (79), our findings provide direct quantitative evidence of how this bias impacts downstream ecological interpretation and viral community structure in the human faecal virome. Although Wang et al. (21) reported similar diversity and composition between MDA viromes and non-MDA viromes, the universal use of a 3,000 bp contig cutoff likely excluded most ssDNA viruses, masking MDA biases. Our beta-diversity comparisons (Fig. 2c) confirm that such biases can be computationally addressed but not fully corrected for. Of course, as with all analyses presented here, the emphasis on comparisons across methods from the same sample does not allow for generalizability across individuals, but we have demonstrated that MDA can result in extreme bias.

Our findings suggest that while MDA viromes may still resolve broad compositional patterns, they remain inappropriate for detailed ecological and functional analyses, as they cannot fully compensate for the loss of diversity or sequencing depth, nor can the simple approach of removing ssDNA viral sequences. Thus, while MDA viromes may still be informative for detecting broad-scale differences in community structure, especially in samples with low viromic DNA yields and/or in re-analyses of datasets already amplified with MDA, our results suggest that using MDA in future viromic studies is difficult to justify. Where its use is unavoidable, computational corrections may help.

### Metagenome-derived viral communities differ from viromes in predicted lifestyles, host associations, and genome annotations

It would be logical to presume that metagenomes would recover more temperate phages (viruses capable of both lytic replication and lysogenic integration into host genomes) than viromes that target free viral particles (80,81), and here we tested that assumption and explored the differences in viral communities recovered from metagenomes and viromes. We first used BACPHLIP (62) to predict the lifestyle (temperate, virulent, or unclassified) of each vOTU, which uses a random forest-based classifier trained on the presence or absence of lysogeny-associated protein domains. Consistent with previous reports of widespread lysogeny in the human gut (79), putative temperate phages were more abundant than those predicted to be virulent across all processing methods, including viromes (Fig. 3a), suggesting that many virions recovered from viromes retain the capacity for lysogenic infection. A substantially smaller proportion of the vOTUs from MDA viromes could be assigned a putative viral lifestyle, likely due to the dominance of *Microviridae*, which are expected to be challenging for viral lifestyle prediction algorithms. Specifically, while some *Microviridae* can integrate into host genomes via co-option of host cell XerC/XerD recombinases (82), they lack phage-borne integrases or transposases, which may prevent BACPHLIP and other prediction tools from predicting a viral lifestyle. To assess the prevalence of integration specifically, we next examined the percentage of temperate phages identified as integrated prophages by geNomad, which uses a conditional random field model to detect genomic regions enriched with viral markers and flanked by host chromosomal features (45). This analysis mirrored trends from the BACPHLIP results, i.e., a lower proportion of prophages were detected in MDA viromes and a much higher proportion in metagenomes (Fig. 3b; ANOVA, F_3,8_ = 251.710, p < 0.00001). The detection of putative integrated prophage sequences in viromes may reflect a combination of factors, including: (1) excised host DNA packaged into virions, (2) false-positive prophage boundary predictions, (3) prophages in ultra-small bacteria that may have passed through the 0.2 µm filter, and (4) the presence of residual host DNA/cellular contamination in VLP-concentrates (though we note that, for substantial contributions from free DNA, we would expect a difference between DNase-treated and untreated viromes, which was not observed). Overall, results suggest a greater proportion of integrated prophages recovered from the metagenomes compared to all viromes.

**Figure 3.**
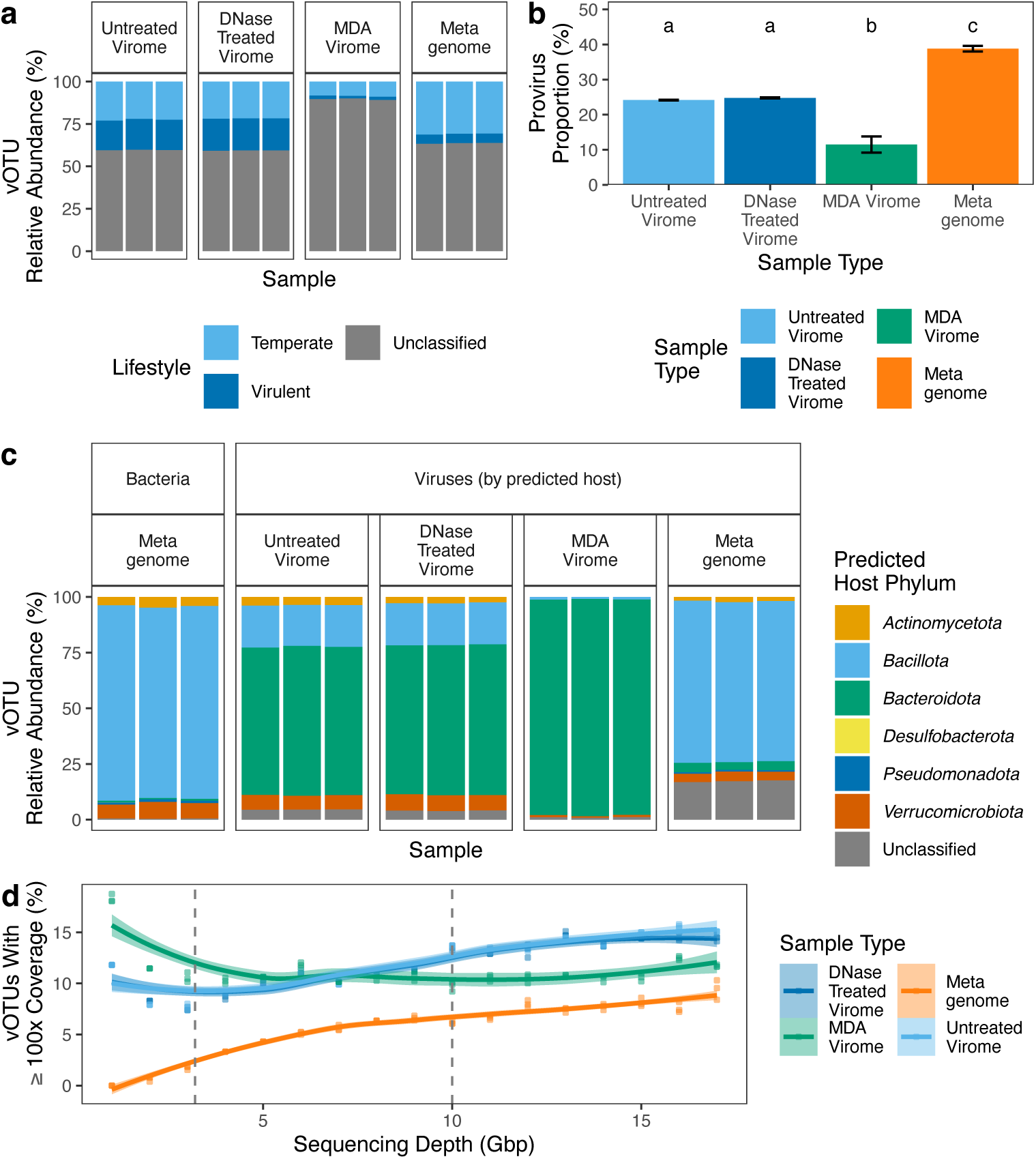
Method-dependent differences in viral lifestyle, host associations and sequencing coverage depth. (a) Relative abundance of predicted viral lifestyles (temperate, virulent, or unclassified) across methods, based on classification by BACPHLIP (62). (b) Percentage of temperate vOTUs predicted to be prophages by geNomad (45). Bars show mean ± standard deviation, letters indicate significant differences between groups (Tukey HSD, p_adjusted_<0.05). (c) vOTU relative abundances by predicted host phyla, according to processing method (left) and metagenome-derived prokaryotic community composition (far right).

We next explored whether predicted prokaryotic host taxa were similarly distributed in the viral communities recovered from each processing method and how these predicted host distributions compared to the prokaryotic community compositions recovered from the metagenomes. Hosts for each vOTU were predicted using iPHoP, and bacterial community profiles were characterised by SingleM (38,63) (Fig. 3c). Bacterial communities were dominated by *Bacillota* (formerly *Firmicutes* (83)), a pattern reflected in the predicted host associations of vOTUs identified in the metagenomes but not the viromes. This is consistent with the observed greater proportion of predicted prophages in the metagenomes compared to viromes, as viral and host abundance should be more highly correlated in the metagenomes, where they are more often coming from the same chromosome. Although viromes and metagenomes generally yielded comparable numbers of vOTUs predicted to infect each bacterial phylum, the relative abundance distributions of these vOTUs differed substantially across processing methods. For example, phages predicted to infect the phylum *Bacillota* comprised 72% of the viral community by relative abundance in metagenomes but only 19% in DNase-treated viromes, despite comparable proportions by vOTU counts (60% and 51%, respectively). In contrast, viruses predicted to infect the phylum *Bacteroidota* dominated all viromes (67% relative abundance of non-MDA viromes, 97% of MDA viromes) but were rare in metagenome viral communities (4%) and virtually absent in the prokaryotic communities (0.7%). Together, these results suggest that different processing methods could lead to wildly different interpretations of infection dynamics, since viral relative abundances differed dramatically between viromes and metagenomes when grouped according to predicted hosts.

We compared profiles of functionally annotated viral genes, including putative auxiliary metabolic genes (AMGs) (1), across processing methods. To more fully capture viral gene content, analyses were conducted at the assembled contig (as opposed to vOTU) level, after meeting viral prediction thresholds (see Methods). Using Pharokka (61), we found that 0.9-1.3% of genes across all viral contigs were annotated as moron elements (transferable elements within phage genomes (84)), accessory metabolic genes, or were involved in host takeover, with no significant differences across viromic methods (Tukey HSD, p > 0.05). However, this category of gene was ∼33% more frequently detected in metagenome-derived viral contigs compared to those from DNase-treated viromes (Supplementary Fig. S6a), suggesting greater prevalence in host-associated viral genomes (e.g., in integrated or actively replicating viruses), as previously suggested (1), and/or more false-positive viral predictions or errors predicting prophage boundaries in metagenomes. Accessory genes were rare overall, with significant differences only between viromes and metagenomes. This overrepresentation in metagenome-derived viral contigs may reflect false-positive detection of host-derived sequences rather than true viral gene content, and underscores the need for rigorous quality control when analysing putative AMGs (85,86).

### Methodological Impact on Viral Prediction Confidence and Viral Recovery at Increasing Sequencing Depths

To assess whether vOTU quality might contribute to observed differences between viromes and metagenomes, we next examined viral prediction confidence scores across processing methods. geNomad confidence scores differed significantly (Fig. 4a, ANOVA, F_3,8_ = 47.5, p = 1.92 × 10⁻⁵), but the mean shift between processing methods was only a modest shift of 0.8–1.7%. Cumulative frequency diagrams (Fig. 4a) show that metagenomes contain a greater proportion of lower-confidence viral contigs, underscoring the need for additional caution when analysing metagenome-derived viral sequences, as a small shift in confidence score distribution can result in the inclusion of tens to hundreds of lower-quality viral sequences. Conversely, 2,261 contigs were excluded through length-filtering but had viral confidence scores >0.95. These shorter, high-confidence contigs may still be useful in specific contexts, but this choice must be clearly justified to avoid inclusion of false positives.

**Figure 4.**
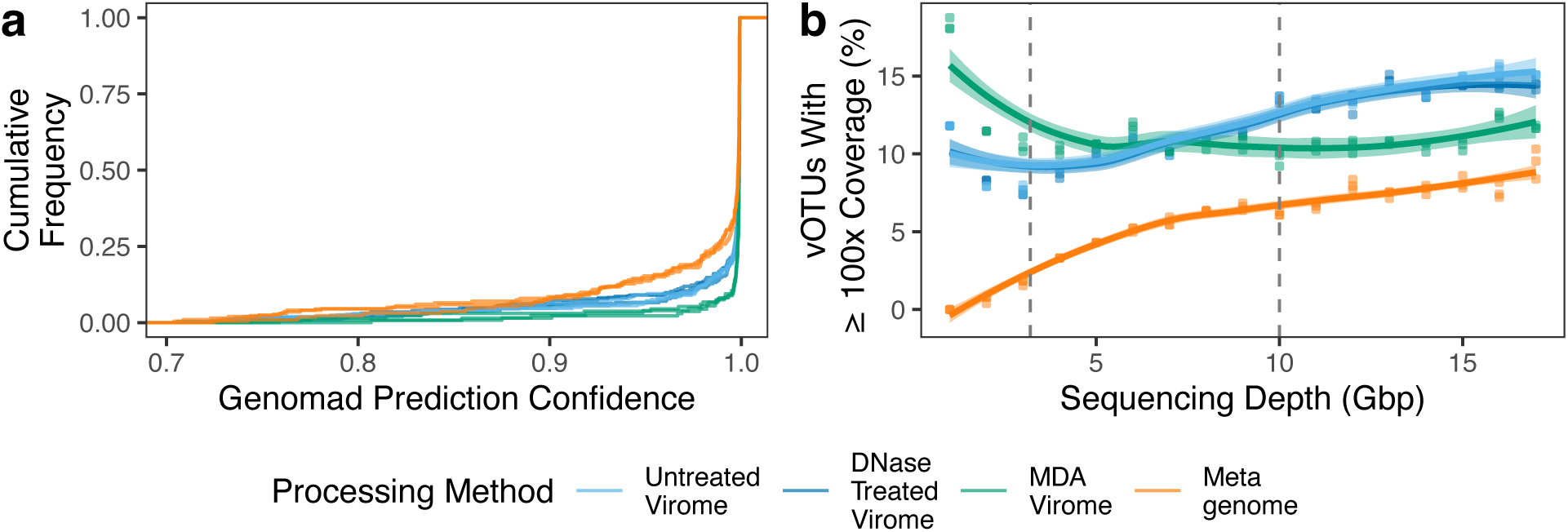
Impact of processing method on viral prediction and high-coverage viral contigs. (a) Cumulative frequency diagram of geNomad prediction confidence scores of viral contigs filtered for length ≥10 Kbp or between 1 and 10 Kbp and geNomad assigned phylum = Monodnaviria. (b) Number of vOTUs with ≥100 × coverage across subsampled sequencing depths (Gbp) for each method. Lines and shaded areas indicate fitted LOESS curves with 95% confidence intervals. Dashed vertical lines indicate commonly used viral dataset sequencing targets, 3.2 Gbp (56) and 10 Gbp (69,87–89).

One presumed advantage of viromes over metagenomes is improved recovery of strain-level microdiversity, which requires high coverage depth (90,91). To compare coverage depth for vOTUs across processing methods and sequencing effort, we subsampled the raw reads from each library in 1 Gbp increments between 1 and 17 Gbp and reprocessed each subset independently. Metagenomes consistently had the lowest proportion of vOTUs with ≥100 × average coverage (Fig. 4b), indicating, as expected, that viromic methods offer better access to sufficient coverage depth for strain-level analyses. At depths <7 Gbp, MDA viromes outperformed non-MDA viromes in recovering high-coverage contigs, reflecting lower viral diversity in MDA viromes and leading to higher coverage. However, this trend was reversed at higher sequencing efforts, with non-MDA viromes consistently capturing vOTUs with higher coverage. In non-MDA viromes, high-coverage vOTUs plateaued at ≤14% of all vOTUs after 12.5 Gbp, suggesting that per-sample strain-level analyses would likely only be possible for high-abundance vOTUs. Our results further confirm that non-MDA viromes outperform both metagenomes and MDA viromes in generating high-coverage vOTUs suitable for detailed strain-level analyses.

### The majority of recovered vOTUs were absent from existing human gut virome databases

The extent to which existing reference databases capture gut virome diversity is unknown, limiting our understanding of the human gut virosphere. To assess the novelty of viruses recovered by this study according to processing method, we clustered all 2,480 recovered viral contigs with the Unified Human Gut Virome catalogue (UHGV, https://github.com/snayfach/UHGV - accessed June 12^th^ 2025), a non-redundant set of 168,536 vOTUs from 12 large-scale datasets (31,49,51–60). Species-level clustering yielded 603 vOTU clusters containing at least one viral contig from this study, compared to 605 vOTUs formed from our study alone. Only 35% (214 of 603) of clusters included UHGV sequences, meaning 65% of our recovered vOTUs were not present in the reference database (Supplementary Fig. S7). Read-mapping to the UHGV database identified a further 248 UHGV vOTUs that were not assembled *de novo* here (Supplementary Fig. S7c), with non-MDA viromes detecting 17% more than metagenomes and 130% more than MDA viromes (Supplementary Fig. S7d). These mapped-only vOTUs had similar coverage (Supplementary Fig. S7e) but were, on average, 50% shorter in length than clustered vOTUs (Supplementary Fig. S7f). These vOTUs may represent viral genomes with intrinsic features, such as high microdiversity or repetitive elements, that hinder complete assembly. Consistent with previous studies (2), our results indicate that a substantial fraction of human gut viral diversity remains uncharacterised and that non-MDA viromes best recover low-abundance viruses, both novel and reference-mapped but unassembled vOTUs.

### Conclusions and outlook

We demonstrate how methodological choices in faecal DNA preparation can shape the composition of the recovered gut viral community and downstream interpretations. MDA introduced a striking bias toward *Microviridae*, consistent with prior studies of ssDNA overamplification (23–25,92–95). Although computationally filtering out short ssDNA contigs can reduce this distortion, MDA remains unsuitable for more detailed ecological comparisons. Viromes had higher viral richness, fewer putative temperate phages (viruses capable of switching between lytic replication and lysogeny), and fewer integrated prophages than metagenomes, emphasising the need to align methodological choices with study objectives.

We also showed that differences in viral community composition due to preparation methods can propagate through downstream analyses beyond community diversity metrics, with the potential for substantially influencing ecological and functional conclusions. We applied a suite of commonly used analytical approaches to examine how methodological choice affected interpretation, resulting in an analytical framework that could be useful for future methodological comparisons. For example, MDA viromes yielding fewer lifestyle predictions due to *Microviridae* dominance, and metagenomes showed substantially different predicted host compositions of the viral community at the phylum level compared to both non-MDA and MDA viromes. Had each method been used in separate studies, these differences could have led to conflicting interpretations, as they produce methodological artefacts that can be difficult to recognise in the literature without expert technical evaluation of study methodologies (19,96,97).

Despite the study’s limited scope, the consistency across replicates and the magnitude of the observed effects emphasise the importance of understanding the sources of methodological bias and potential mitigation methods. The more refined methods described herein will have significant value when applied to larger-scale human studies that explore interindividual variability of the gut virome and associations of the virome with health phenotypes.

Our findings complement and extend prior gut virome methods comparisons, which have often focused on individual steps, such as viral enrichment, DNA extraction, and sequencing library construction (15,19,21,30,98,99). No single method can capture all aspects of gut viral diversity, but by understanding the specific biases and strengths of each technique, researchers can make more informed methodological choices. As larger-scale, more integrative and multi-omic studies become more prevalent, careful method selection will become increasingly essential for generating robust insights into the role of the gut virome in human health and disease.

## Acknowledgments

Elements of the graphical abstract and Supplementary Fig. S1 were produced in BioRender. The sequencing was carried by the DNA Technologies and Expression Analysis Core at the UC Davis Genome Center, supported by NIH Shared Instrumentation Grant 1S10OD010786-01. The authors also thank other members of the Emerson Lab for helpful discussions and general feedback.

## Authors’ contributions

LSH: conceptualisation, data curation, formal analysis, writing – original draft preparation, visualization

TAK: resources, project administration, writing – review and editing

SHA: resources, project administration, writing – review and editing MRA: resources, project administration, writing – review and editing

MO: conceptualisation, formal analysis, visualization, writing – review and editing

JBE: conceptualisation, wet-lab methodology, investigation, writing – review and editing, supervision, funding acquisition, project administration

## Funding

This research was supported by the NIH Common Fund (Human Virome Program), Award # U01DE034198. LSH was also partially supported by the U.S. Department of Energy (DOE), Office of Science, Office of Biological and Environmental Research (BER), Genomic Science Program, award number DE-SC0023127 (grant to JBE, grant PI Sydney Glassman). Research on the human microbiome by Drs. Adams, Knotts, and Ali is funded, in part, by NIH-NIDDK 1R01DK137173-01A1.

## Data and Code Availability

All raw sequencing data have been deposited with the SRA under BioProject PRJNA1331857 (SRR35523994 - SRR35524005). As sequencing data deposition requires the removal of human metagenomic reads, human reads were scrubbed from the raw read files with the NCBI’s Human Read Removal Tool (https://hub.docker.com/r/ncbi/sra-human-scrubber) prior to submission. Human read count tables are archived in the GitHub and Zenodo repositories. All filtered viral sequences were deposited in GenBank and in the GitHub/Zenodo repositories for the manuscript. All R scripts involved in the processing and analysis of these data, plus summary tables used as input, have been deposited on GitHub (https://github.com/LSHillary/FecalViromeOptimisation) and archived on Zenodo (DOI: 10.5281/zenodo.17527407).

## Competing interests

S.H. Adams is founder and principal of XenoMed, LLC (dba XenoMet), which is focused on research and discovery in microbial metabolism. XenoMed had no part in the research design, funding, results or writing of the manuscript.

## List of Abbreviations

AMG: Auxiliary Metabolic Gene
ANI: Average Nucleotide Identity
dsDNA: double-stranded DNA
MDA: Multiple Displacement Amplification
ssDNA: single-stranded DNA
VLP: Virus-Like-Particle
vOTU: viral Operational Taxonomic Unit

