## Supplementary Information for "DNA extraction and virome processing methods strongly influence recovered human gut viral community characteristics"

#### Supplementary Tables

**Supplementary Table S1 – Anonymised donor metadata**

| Attribute | Value |
|---|---|
| Sex | Female |
| Age | 55 |
| Exam Date | 2019/05/06 |
| Lab Date | 2019/05/06 |
| Height | 1.65 m |
| Weight | 116 Kg |
| BMI | 42.6 |

**s**

**Supplementary Table S2 – Virome and metagenome DNA yields**

| Sample | Sample Description | MDA | DNA Concentration (ng/ µL) |
|---|---|---|---|
| **F3_S19** | DNase-Treated Virome | FALSE | 1.10 |
| **F4_S20** | DNase-Treated Virome | FALSE | 1.12 |
| **F5_S21** | DNase-Treated Virome | FALSE | 1.29 |
| **F6_S22** | Untreated Virome | FALSE | 2.39 |
| **F7_S23** | Untreated Virome | FALSE | 3.01 |
| **F8_S24** | Untreated Virome | FALSE | 2.23 |
| **M1_S25** | Metagenome | FALSE | 143.5 |
| **M2_S26** | Metagenome | FALSE | 54.5 |
| **M3_S27** | Metagenome | FALSE | 52.0 |
| **GP3_S28** | MDA Amplified Virome | TRUE | 47.4 |
| **GP4_S29** | MDA Amplified Virome | TRUE | 54.0 |
| **GP5_S30** | MDA Amplified Virome | TRUE | 60.0 |

**Supplementary Table S3 – Settings of bioinformatics data processing steps and R packages used in this study.** Data processing, analysis and visualization scripts can be found in this study's GitHub/ Zenodo repository (https:// https://github.com/LSHillary/FecalViromeOptimisation and DOI: 10.5281/zenodo.17527407).

| Step | Program | Settings | Reference |
|---|---|---|---|
| Raw read inspection | FastQC v0.12.1 | Default settings | (Andrews, 2010) |
| Combining FastQC reports | MultiQC v1.14 | Default settings | (Ewels et al., 2016) |
| Read filtering and quality trimming | BBDuk v39.1 | ref=adapters,phix ktrim=r k=23 mink=11 hdist=1 tpe tbo qtrim=r trimq=10 maxns=3 maq=3 minlen=50 mlf=0.333 | (Bushnell, 2018) |
| Error correction | Tadpole | mode=correct ecc=t prefilter=1 | (Bushnell, 2018) |
| Read PCR duplicate removal | Clumpify | dedupe subs=0 passes=2 | (Bushnell, 2018) |
| K-mer frequency analysis | Khmer v2.1.1 | -k 31 (trim-low-abund.py -V -C 2) | (Crusoe et al., 2015) |
| Ribosomal rRNA read identification in error corrected reads | SortMeRNA v4.3.6 | --paired_in --num_alignments 1 --out2 –fastx -v (Database = smr_v4.3_sensitive_db) | (Kopylova et al., 2012) |
| Raw read taxonomic profiling | SingleM v0.18.3 | singlem pipe (default settings) | (Woodcroft et al., 2024) |
| Assembly of raw reads into contigs | MEGAHIT v1.2.9 | --k-min 27 --min-contig-len 1000 --presets meta-large | (Li et al., 2015) |
| Assembly summary statistics | Quast v5.2.0 | quast.py (default settings) | (Mikheenko et al., 2018) |
| Identification of viral contigs | GeNomad v1.7.0. (Database v1.4) | end-to-end --cleanup --enable-score-calibration | (Camargo et al., 2024) |
| Clustering of viral contigs into viral operational taxonomic units | MegaBLAST v2.14.0<br>Custom scripts derived from CheckV, provided in this study's GitHub | See this study's GitHub repository. Pipeline settings:<br>leiden_resolution=1.0<br>min_ani=0.95<br>min_cov=0.85<br>blast_max_evalue=1e-5 | (Camargo et al., 2023; Morgulis et al., 2008; Nayfach et al., 2020) |
| Mapping error corrected reads | Minimap2 v2.26 | -axsr | (Li, 2018; Nurk et al., 2022) |

| | | | |
|---|---|---|---|
| to the human genome/ viral contigs | Refseq assembly GCF_009914755.1 | | |
| Conversion of SAM files to sorted BAM files | Samtools v1.17 | samtools view -u \| samtools sort | (Li et al., 2009) |
| Calculate mapped read counts and tpm values | CoverM v0.6.1 | For tpm:<br>coverm contig --methods mean trimmed_mean covered_bases variance rpkm tpm --min-read-percent-identity 90 --min-covered-fraction 75 --output-format dense<br><br>For read counts:<br>--methods mean trimmed_mean count | (Aroney et al., 2025) |
| Subsampling of raw reads for assessing effects of read depth on viral genome recovery | Seqktk v1.4-r122 | seqtk sample -s 100 <number of reads> | (https://github.com/lh3/seqtk) |
| Host Prediction | iPhop v1.3.2 | iphop predict (Database = Aug_2023_pub_rw) | (Roux et al., 2023) |
| Viral protein annotation | Pharokka v1.7.1 | pharokka_proteins.py -f (Database = v1.4.0) | (Bouras et al., 2023) |
| Viral lifestyle prediction | BACPHLIP v0.9.6 | --multi_fasta -f | (Hockenberry and Wilke, 2021) |
| Data analysis and visualization | R v4.4.0 | See GitHub repository for detailed R scripts and functions used in this study | (R Core Team, 2024) |
| | RStudio v2024.09.1-394 | See GitHub repository for detailed R scripts and functions used in this study | (Posit team, 2024) |
| Visualizing Venn diagrams | ggVennDiagram v1.5.2 | | |
| Data visualization | ggpubr v0.6.0 | | (Kassambara, 2020) |
| Assessing statistically significant differences | multcompView v0.1.10 | | (Graves et al., 2015) |
| Data visualization | scales v1.3.0 | | (Wickham et al., 2011) |

| | | | |
|---|---|---|---|
| Data analysis pipeline management | targets v1.10 .0 | | (Landau, 2021) |
| Data wrangling and visualization | tidyverse v2.0.0 | | (Wickham et al., 2019) |
| PCoA production | vegan 2.6.8 | | (Oksanen et al., 2019) |

**Supplementary Table S4 – Percentage of unique and shared vOTUs by sample type assessed by read mapping.** DNase-treated and untreated viromes were summed for these calculations as only 12 vOTUs (2.6%) were not shared between these two methods.

| Sample type | Total mapped vOTUs | Unique vOTUs | Shared with both other methods |
|---|---|---|---|
| DNase treated/ untreated viromes | 458 | 48 (10.4%) | 150 (32.8%) |
| MDA-viromes | 293 | 24 (8.2%) | 150 (51.2 %) |
| Metagenomes | 418 | 121 (28.9%) | 150 (35.9 %) |

### Supplemental Figures

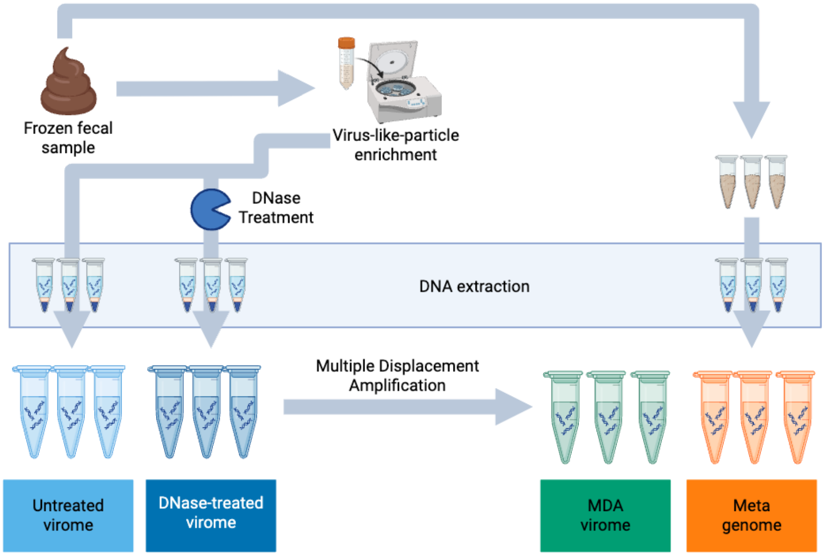


**Supplementary Figure S1 – Experimental design.** A single frozen fecal sample was subsampled for total faecal DNA extraction and virus particle enrichment. The VLP concentrate was split and half the volume was treated with DNase prior to DNA extraction. An aliquot of DNase treated VLP DNA extract was used for multiple displacement amplification. Created in BioRender. Hillary, L. (2025) https://BioRender.com/s4h7hc9

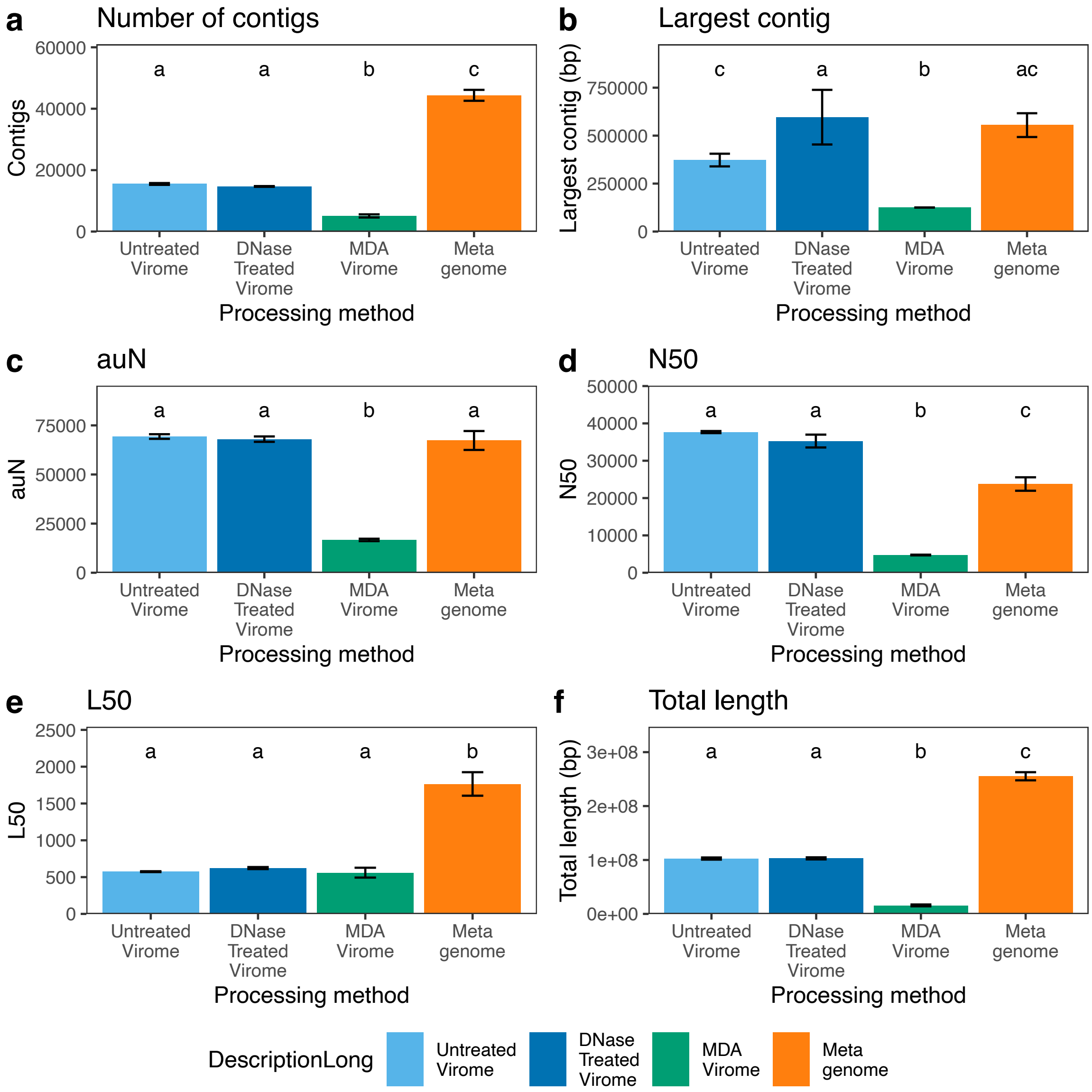


**Supplementary Figure S2 – Summary contig statistics** showing the distribution of (a) the number of contigs, (b) longest contig, (c) auN (area under the Nx curve), (d) N50 (length of the shortest contig needed to be included to cover 50% of the total assembly length from contigs ordered from longest to shortest) , (e) L50 (rank of the N50 contig from longest to shortest) and (f) total assembly length. Error bars represent mean ± standard deviation, and letters indicate groupings based on statistically significant differences ($p < 0.05$) from Tukey HSD tests. Full details of the test statistics can be found in supplementary tables S5 and S6.

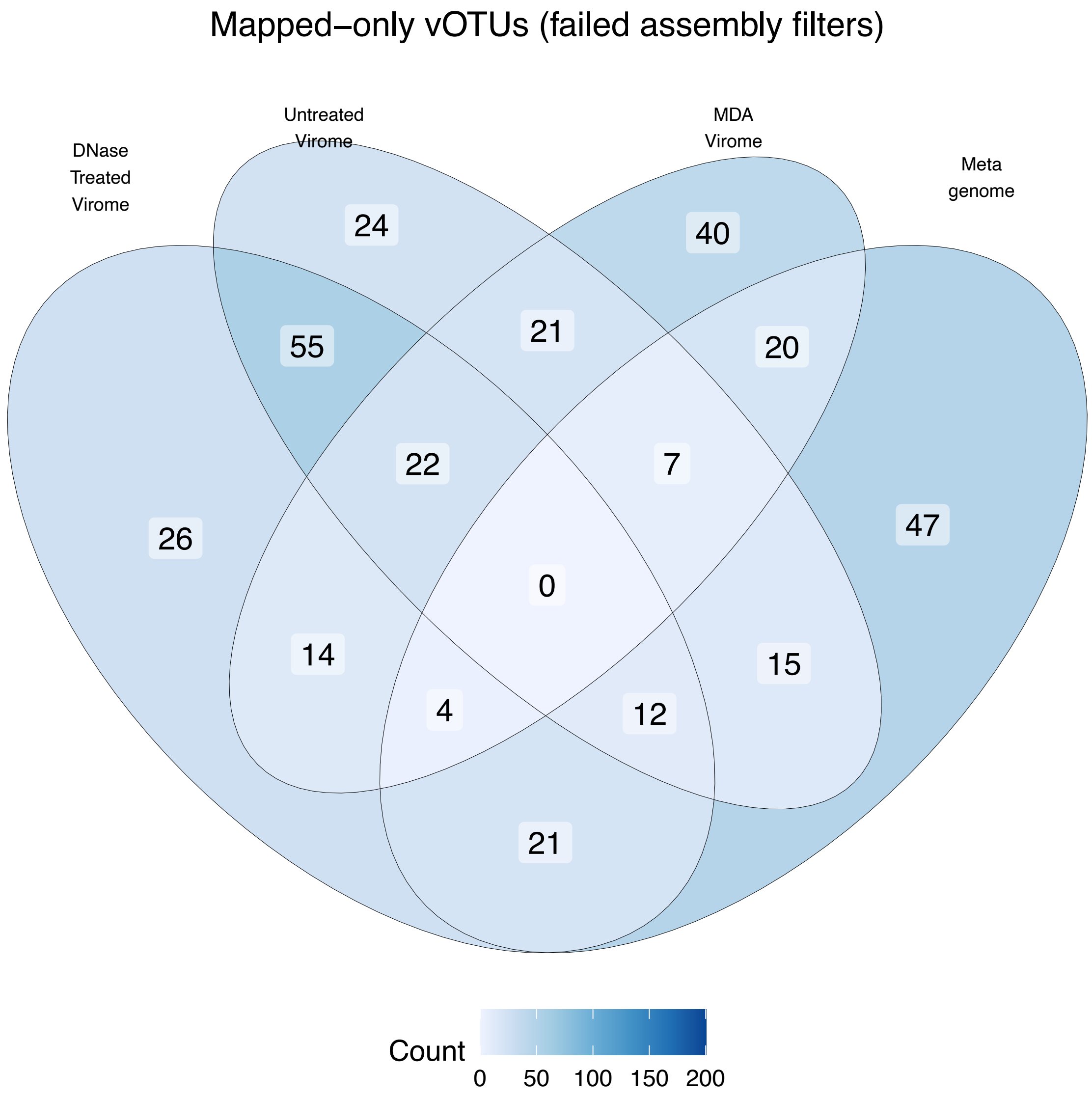


**Supplementary Figure S3** – Venn diagram of vOTUs that were only detected by mapping and not assembly across sample types.

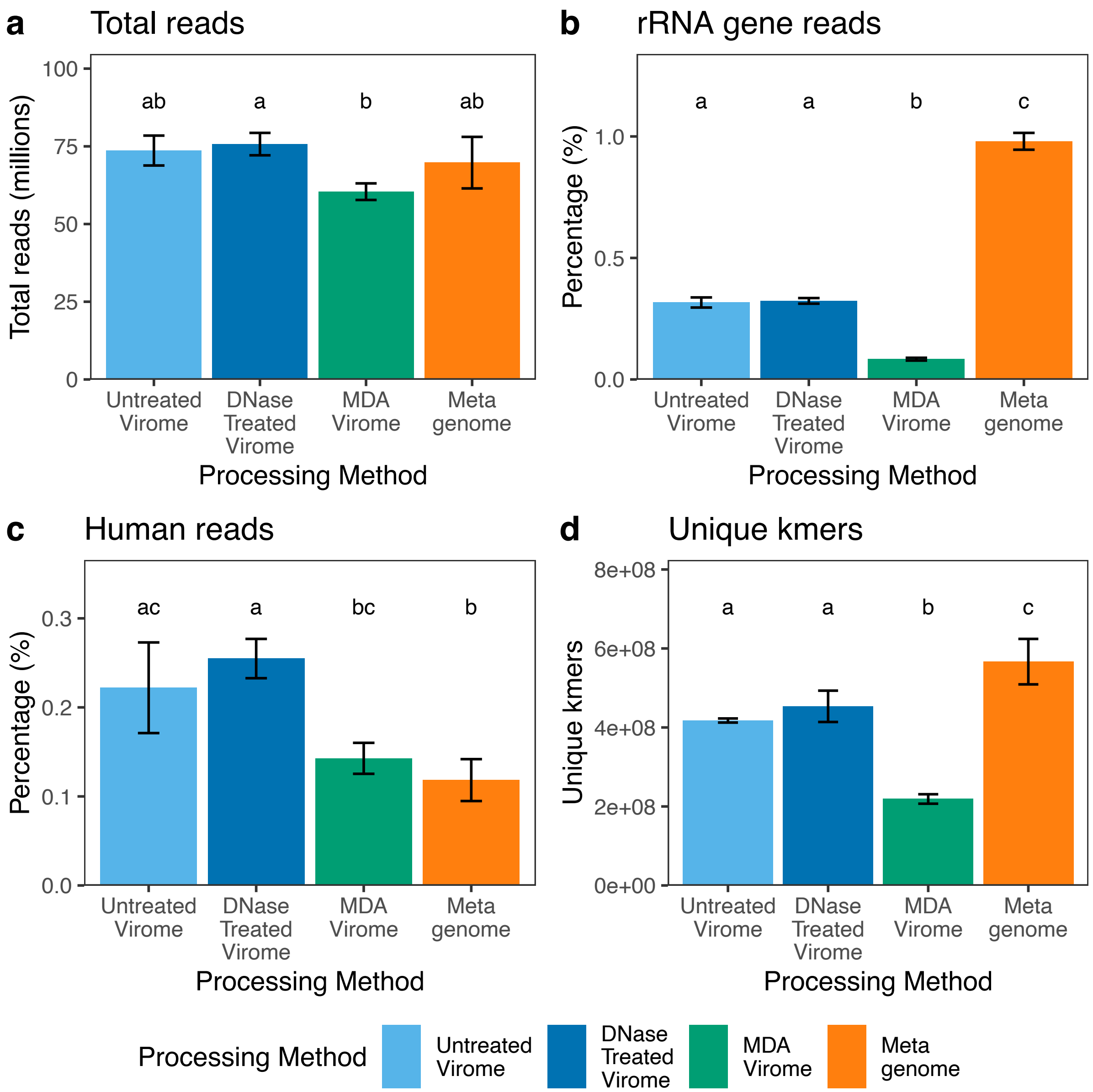


**Supplementary Figure S4 – Read summary statistics.** (a) total raw reads, (b) rRNA gene reads, (c) human reads, and (d) unique k-mer counts for each processing method. Letters represent compact letter display groupings based on Tukey HSD post-hoc tests ($p < 0.05$). Full details of the test statistics can be found in supplementary tables S5 and S6.

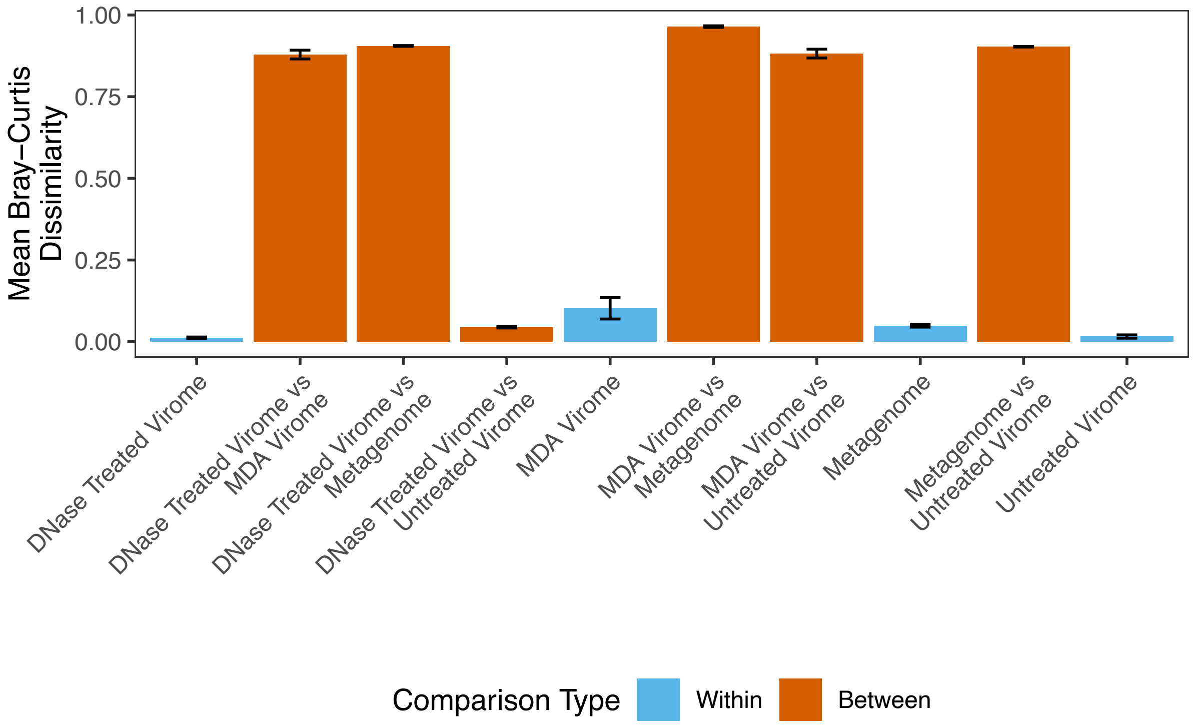


**Supplementary Figure S5 – Mean Bray-Curtis dissimilarities within and between virome and metagenome sample types.** Bars represent the mean Bray-Curtis dissimilarity ± standard deviation for each sample type comparison. “Within” comparisons reflect mean dissimilarity between replicates of the same sample type, while “Between” comparisons represent dissimilarity between replicates of different types. Although some vOTUs were shared across methods, between-group dissimilarities were consistently high, with the exception of DNase-treated and untreated viromes (0.04).

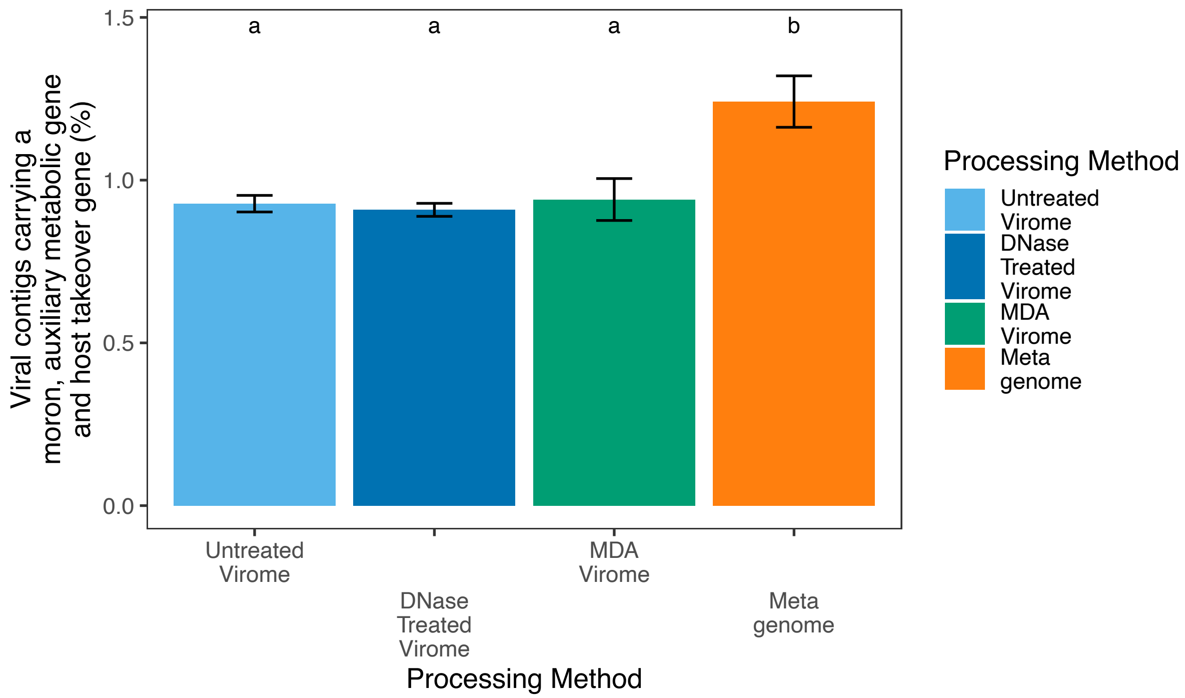


**Supplementary Figure S6 – Viral contig annotations from each sample type.** Values were calculated from annotations of assembled viral contigs from individual samples. Metagenomes had significantly more moron/ AMG/ host takeover genes than viromes. Error bars represent mean ± standard deviation, and letters indicate groupings based on statistically significant differences ($p < 0.05$) from Tukey HSD tests. Full details of the test statistics can be found in supplementary tables S5 and S6.

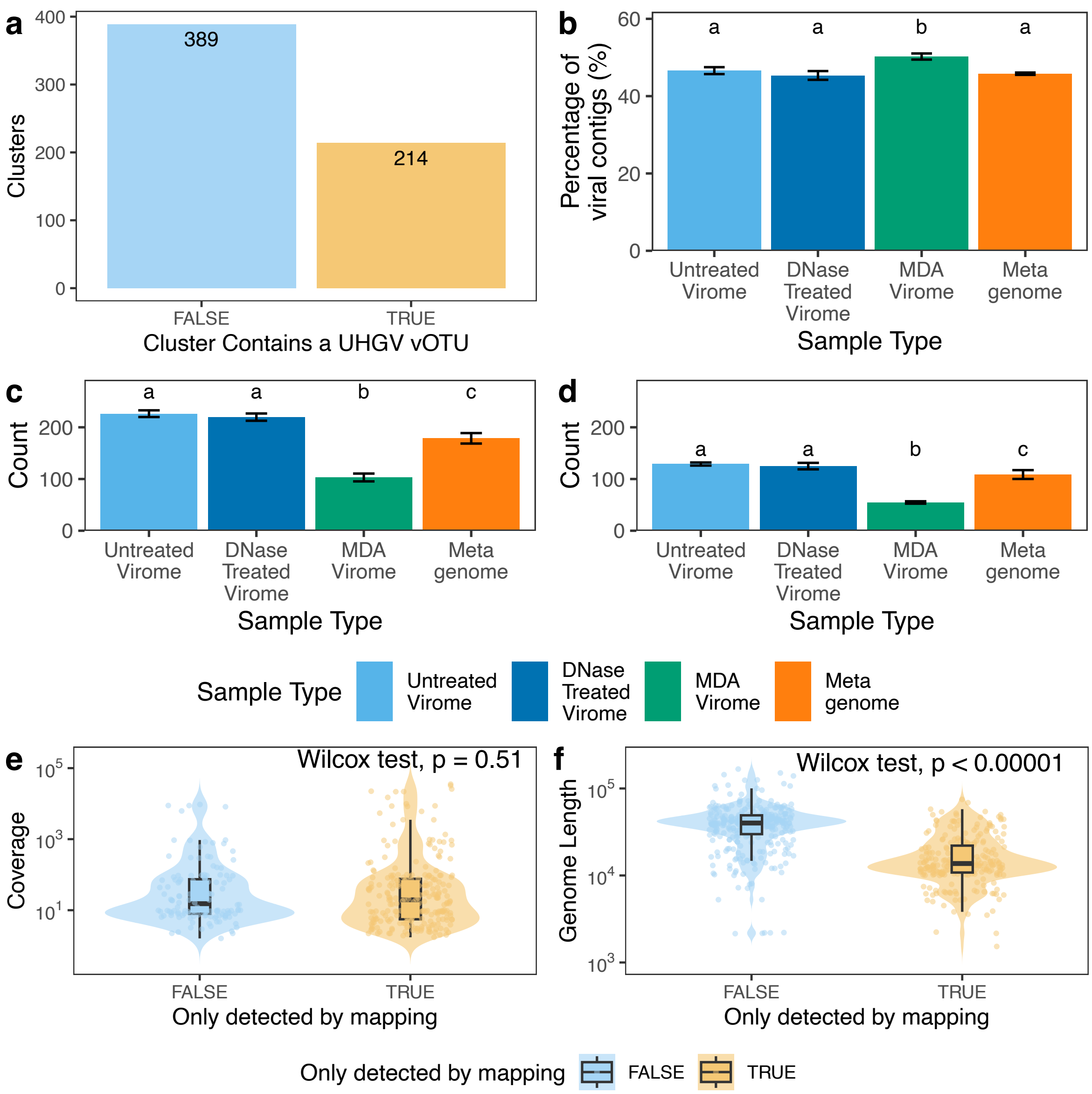


**Figure S7 – Comparison of this study's vOTUs with those found in the Unified Human Gut Virome Catalogue (UHGV)** (a) Number of clusters containing at least one viral contig from this study and one vOTU from the UHGV. (b) The percentage of viral contigs assembled from each sample type to cluster with a UHGV vOTU. (c) The number of UHGV vOTUs detected by read mapping across different sample types and (d) those vOTUs only detected by mapping. Clustering and read mapping was performed in the same way as with the analysis of contigs/ reads from this study. Bars show mean ± standard deviation, letters indicate significant differences between groups (Tukey HSD, $p<0.05$). Full details of the test statistics can be found in supplementary tables S5 and S6. (e) The distribution of coverage values of UHGV contigs identified in samples from this study based on clustering and mapping, or just mapping and (f) the distribution of genome length of UHGV contigs identified in samples from this study based on clustering and mapping, or just mapping.

#### Supplementary Information References

Andrews, S., 2010. FastQC: A Quality Control Tool for High Throughput Sequence Data.

Aroney, S.T.N., Newell, R.J.P., Nissen, J.N., Camargo, A.P., Tyson, G.W., Woodcroft, B.J., 2025. CoverM: read alignment statistics for metagenomics. Bioinformatics 41, btaf147. https://doi.org/10.1093/bioinformatics/btaf147

Bouras, G., Nepal, R., Houtak, G., Psaltis, A.J., Wormald, P.-J., Vreugde, S., 2023. Pharokka: a fast scalable bacteriophage annotation tool. Bioinformatics 39, btac776. https://doi.org/10.1093/bioinformatics/btac776

Bushnell, B., 2018. BBTools: a suite of fast, multithreaded bioinformatics tools designed for analysis of DNA and RNA sequence data.

Camargo, A.P., Nayfach, S., Chen, I.-M.A., Palaniappan, K., Ratner, A., Chu, K., Ritter, S.J., Reddy, T.B.K., Mukherjee, S., Schulz, F., Call, L., Neches, R.Y., Woyke, T., Ivanova, N.N., Eloe-Fadrosh, E.A., Kyrpides, N.C., Roux, S., 2023. IMG/VR v4: an expanded database of uncultivated virus genomes within a framework of extensive functional, taxonomic, and ecological metadata. Nucleic Acids Res. 51, D733–D743. https://doi.org/10.1093/nar/gkac1037

Camargo, A.P., Roux, S., Schulz, F., Babinski, M., Xu, Y., Hu, B., Chain, P.S.G., Nayfach, S., Kyrpides, N.C., 2024. Identification of mobile genetic elements with geNomad. Nat. Biotechnol. 42, 1303–1312. https://doi.org/10.1038/s41587-023-01953-y

Crusoe, M.R., Alameldin, H.F., Awad, S., Boucher, E., Caldwell, A., Cartwright, R., Charbonneau, A., Constantinides, B., Edvenson, G., Fay, S., Fenton, J., Fenzl, T., Fish, J., Garcia-Gutierrez, L., Garland, P., Gluck, J., González, I., Guermond, S., Guo, J., Gupta, A., Herr, J.R., Howe, A., Hyer, A., Härpfer, A., Irber, L., Kidd, R., Lin, D., Lippi, J., Mansour, T., McA'Nulty, P., McDonald, E., Mizzi, J., Murray, K.D., Nahum, J.R., Nanlohy, K., Nederbragt, A.J., Ortiz-Zuazaga, H., Ory, J., Pell, J., Pepe-Ranney, C., Russ, Z.N., Schwarz, E., Scott, C., Seaman, J., Sievert, S., Simpson, J., Skennerton, C.T., Spencer, J., Srinivasan, R., Standage, D., Stapleton, J.A., Steinman, S.R., Stein, J., Taylor, B., Trimble, W., Wiencko, H.L., Wright, M., Wyss, B., Zhang, Q., Zyme, E., Brown, C.T., 2015. The khmer software package: enabling efficient nucleotide sequence analysis. https://doi.org/10.12688/f1000research.6924.1

Ewels, P., Magnusson, M., Lundin, S., Käller, M., 2016. MultiQC: summarize analysis results for multiple tools and samples in a single report. Bioinformatics 32, 3047–3048. https://doi.org/10.1093/bioinformatics/btw354

Graves, S., Piepho, H.-P., Selzer, M.L., 2015. Package 'multcompView.' Vis. Paired Comp. 451, 452.

Hockenberry, A.J., Wilke, C.O., 2021. BACPHLIP: predicting bacteriophage lifestyle from conserved protein domains. PeerJ 9, e11396. https://doi.org/10.7717/peerj.11396

Kassambara, A., 2020. ggpubr: "ggplot2" Based Publication Ready Plots.

Kopylova, E., Noe, L., Touzet, H., 2012. SortMeRNA: fast and accurate filtering of ribosomal RNAs in metatranscriptomic data. Bioinformatics 28, 3211--3217. https://doi.org/10.1093/bioinformatics/bts611

Landau, W.M., 2021. The targets R package: a dynamic Make-like function-oriented pipeline toolkit for reproducibility and high-performance computing. J. Open Source Softw. 6, 2959. https://doi.org/10.21105/joss.02959

Li, D., Liu, C.-M., Luo, R., Sadakane, K., Lam, T.-W., 2015. MEGAHIT: an ultra-fast single-node solution for large and complex metagenomics assembly via succinct de Bruijn graph. Bioinformatics 31, 1674--1676. https://doi.org/10.1093/bioinformatics/btv033

Li, H., 2018. Minimap2: pairwise alignment for nucleotide sequences. Bioinformatics 34, 3094–3100. https://doi.org/10.1093/bioinformatics/bty191

Li, H., Handsaker, B., Wysoker, A., Fennell, T., Ruan, J., Homer, N., Marth, G., Abecasis, G., Durbin, R., 2009. The Sequence Alignment/Map format and SAMtools. Bioinformatics 25, 2078–2079. https://doi.org/10.1093/bioinformatics/btp352

Morgulis, A., Coulouris, G., Raytselis, Y., Madden, T.L., Agarwala, R., Schäffer, A.A., 2008. Database indexing for production MegaBLAST searches. Bioinformatics 24, 1757–1764. https://doi.org/10.1093/bioinformatics/btn322

Nayfach, S., Camargo, A.P., Schulz, F., Eloe-Fadrosh, E., Roux, S., Kyrpides, N.C., 2020. CheckV assesses the quality and completeness of metagenome-assembled viral genomes. Nat. Biotechnol. 39, 578--585. https://doi.org/10.1038/s41587-020-00774-7

Nurk, S., Koren, S., Rhie, A., Rautiainen, M., Bzikadze, A.V., Mikheenko, A., Vollger, M.R., Altemose, N., Uralsky, L., Gershman, A., Aganezov, S., Hoyt, S.J., Diekhans, M., Logsdon, G.A., Alonge, M., Antonarakis, S.E., Borchers, M., Bouffard, G.G., Brooks, S.Y., Caldas, G.V., Chen, N.-C., Cheng, H., Chin, C.-S., Chow, W., de Lima, L.G., Dishuck, P.C., Durbin, R., Dvorkina, T., Fiddes, I.T., Formenti, G., Fulton, R.S., Fungtammasan, A., Garrison, E., Grady, P.G.S., Graves-Lindsay, T.A., Hall, I.M., Hansen, N.F., Hartley, G.A., Haukness, M., Howe, K., Hunkapiller, M.W., Jain, C., Jain, M., Jarvis, E.D., Kerpedjiev, P., Kirsche, M., Kolmogorov, M., Korlach, J., Kremitzki, M., Li, H., Maduro, V.V., Marschall, T., McCartney, A.M., McDaniel, J., Miller, D.E., Mullikin, J.C., Myers, E.W., Olson, N.D., Paten, B., Peluso, P., Pevzner, P.A., Porubsky, D., Potapova, T., Rogaev, E.I., Rosenfeld, J.A., Salzberg, S.L., Schneider, V.A., Sedlazeck, F.J., Shafin, K., Shew, C.J., Shumate, A., Sims, Y., Smit, A.F.A., Soto, D.C., Sović, I., Storer, J.M., Streets, A., Sullivan, B.A., Thibaud-Nissen, F., Torrance, J., Wagner, J., Walenz, B.P., Wenger, A., Wood, J.M.D., Xiao, C., Yan, S.M., Young, A.C., Zarate, S., Surti, U., McCoy, R.C., Dennis, M.Y., Alexandrov, I.A., Gerton, J.L., O'Neill, R.J., Timp, W., Zook, J.M., Schatz, M.C., Eichler, E.E., Miga, K.H., Phillippy, A.M., 2022. The complete sequence of a human genome. Science 376, 44–53. https://doi.org/10.1126/science.abj6987

Oksanen, J., Blanchet, F.G., Friendly, M., Kindt, R., Legendre, P., McGlinn, D., Minchin, P.R., O'Hara, R.B., Simpson, G.L., Solymos, P., Stevens, M.H.H., Szoecs, E., Wagner, H., 2019. vegan: Community Ecology Package.

Posit team, 2024. RStudio: Integrated development environment for R (manual). Posit Software, PBC, Boston, MA.

R Core Team, 2024. R: a language and environment for statistical computing (manual). R Foundation for Statistical Computing, Vienna, Austria.

Roux, S., Camargo, A.P., Coutinho, F.H., Dabdoub, S.M., Dutilh, B.E., Nayfach, S., Tritt, A., 2023. iPHoP: An integrated machine learning framework to maximize host prediction for metagenome-derived viruses of archaea and bacteria. PLOS Biol. 21, e3002083. https://doi.org/10.1371/journal.pbio.3002083

Wickham, H., Averick, M., Bryan, J., Chang, W., McGowan, L.D., François, R., Grolemund, G., Hayes, A., Henry, L., Hester, J., Kuhn, M., Pedersen, T.L., Miller, E., Bache, S.M., Müller, K., Ooms, J., Robinson, D., Seidel, D.P., Spinu, V., Takahashi, K., Vaughan, D., Wilke, C., Woo, K., Yutani, H., 2019. Welcome to the tidyverse. J. Open Source Softw. 4, 1686. https://doi.org/10.21105/joss.01686

Wickham, H., Pedersen, T.L., Seidel, D., 2011. scales: Scale Functions for Visualization. https://doi.org/10.32614/CRAN.package.scales

Woodcroft, B.J., Aroney, S.T.N., Zhao, R., Cunningham, M., Mitchell, J.A.M., Blackall, L., Tyson, G.W., 2024. SingleM and Sandpiper: Robust microbial taxonomic profiles from metagenomic data. https://doi.org/10.1101/2024.01.30.578060
